# Revisiting the regulation of the capsular polysaccharide biosynthesis gene cluster in *Staphylococcus aureus*

**DOI:** 10.1101/614925

**Authors:** Daniela Keinhörster, Andrea Salzer, Alejandra Duque-Jaramillo, Shilpa E. George, Gabriella Marincola, Jean C. Lee, Christopher Weidenmaier, Christiane Wolz

## Abstract

In *Staphylococcus aureus*, the capsular polysaccharide (CP) protects against phagocytosis, but also hinders adherence to endothelial cells and matrix proteins. Its biosynthesis is tightly controlled resulting in a heterogeneous phenotype within a population and CP being mainly detectable in non-growing cells. Capsular biosynthesis genes are encoded by a conserved *capA-P* operon whose expression is driven by an upstream promoter element (P*_cap_*) in front of *capA*. The organization of P*_cap_* is poorly understood, as is the interplay of different regulators that influence the early-Off/late-Heterogeneous *cap* transcription pattern. Here, we demonstrate that P*_cap_* contains a main SigB-dependent promoter. The SigB consensus motif overlaps with a previously described inverted repeat that is crucial for *cap* expression. The essentiality of the inverted repeat is derived from this region acting as a SigB binding site rather than as an operator site for the proposed *cap* activators RbsR and MsaB. Furthermore, P*_cap_* contains an extensive upstream region harboring a weak SigA-dependent promoter and binding sites for the *cap* repressors SaeR, CodY and Rot. We show that heterogeneous CP synthesis is determined by the combination of SigB activity and repressor binding to the upstream region. The direct SigB dependency and the upstream repressors are also sufficient to explain the temporal gene expression pattern at the transcriptional level. However, CP synthesis remains growth phase-dependent even when *capA* transcription is rendered constitutive, suggesting additional post-transcriptional regulatory circuits. Thus, the interference of multiple repressors with SigB-dependent promoter activity as well as post-transcriptional mechanisms ensure the appropriate regulation of CP synthesis.

**Importance:** The majority of bacterial pathogens produce an array of polysaccharides on their surface which are important virulence factors and thus serve as attractive vaccine candidates. However, the synthesis and assembly of these structures is highly variable and tightly regulated at various levels. In the human pathogen *Staphylococcus aureus*, the synthesis of the capsular polysaccharide (CP) is dependent on a complex regulatory network which ensures that CP is produced only in a fraction of stationary phase cells. Here, we determined main regulators that drive the peculiar CP expression pattern. We found that the interplay of the transcriptional repressors Sae, CodY and Rot with the alternative Sigma factor B is responsible for early-Off/late-Heterogeneous expression at the transcriptional level. The data also implicates post-transcriptional mechanisms that may act to avoid conflict in precursor usage by machineries involved in either synthesis of CP or other glycopolymers in growing bacterial cells.

## Introduction

*Staphylococcus aureus* is an opportunistic pathogen that asymptomatically colonizes parts of the human population, thereby increasing the risk of subsequent infections. Its capacity to cause a wide variety of diseases depends on secreted virulence factors as well as cell surface attached proteins and polysaccharides (1–3). The capsular polysaccharide (CP) is one of these cell surface structures playing an important role in *S. aureus* pathogenesis and bacterial evasion of the host immune defences (3, 4). Therefore, it is being discussed as a target for immunotherapy and as a vaccine candidate (5).

CP serotypes 5 and 8 are the two main CP serotypes produced by *S. aureus* strains (6–9). Their structure is highly similar due to the closely related *cap5* and *cap8* gene clusters. These allelic operons consist of 16 genes, *cap5/8A* to *cap5/8P*, whose gene products are involved in CP biosynthesis, O-acetylation, transport and regulation (3, 4, 10). The *cap* operon is thought to be mainly transcribed as a single large 17 kb transcript driven by one principal promoter element (P*_cap_*), located upstream of *capA* (11, 12). While *cap* gene clusters are extremely conserved across *S. aureus* genomes, not all clinically relevant isolates produce CP and acapsular variants may emerge during chronic infections. The loss of CP expression can typically be explained by mutations in any of the *cap* genes essential for CP synthesis or in the promoter region (13, 14). For instance, acapsular strains from the USA300 lineage were attributed to conserved mutations in the *cap5* locus (15). However, this assumption has been recently challenged by the finding that USA300 strains might indeed produce CP during infection (16). In addition to mutations that abolish its production, CP synthesis can also be switched off in response to environmental conditions via a complex regulatory network. Extensive *in vitro* and *in vivo* analyses have shown that *cap* expression is highly sensible to changes in nutrients, pH, CO_2_ and oxygen availability (17–24). Interestingly, CP synthesis was commonly found to be strictly growth phase-dependent and detectable only in the post-exponential growth phase (17, 21, 22, 25, 26). In addition, not all bacteria in a population are CP-positive as revealed by flow cytometry and immunofluorescence of *in vitro* and *in vivo* grown bacteria (21, 22, 27, 28). As only non-encapsulated cells are able to adhere to endothelial cells (21), while CP protects bacterial cells from phagocytosis (29–31), it is likely that CP heterogeneity provides better adaptability of the population as a whole. So far the underlying regulatory mechanisms of this particular expression pattern (early-Off/late-Heterogeneous) are only partially understood.

In general, P*_cap_* activity correlates with CP synthesis, indicating that regulation occurs predominantly at the transcriptional level (12, 22, 32–34). Yet, the data to explain the molecular mechanisms of *cap* regulation are puzzling. The identified transcriptional start site (TSS) is not preceded by a classical Sigma factor consensus sequence (12); instead, several inverted and direct repeats were identified further upstream, amongst which a 10 bp inverted repeat (IR) was shown to be crucial for promoter activity (12). It has been proposed that this IR functions as an operator site for the *cap* activators RbsR and MsaB (35, 36). RbsR also functions as a repressor of the *rbsUDK* operon involved in ribose uptake. While the presence of ribose relieves repression of the *rbsUDK* operon by RbsR, the presence and absence of ribose had no effect on *cap* expression (35). MsaB is described as a transcriptional factor with DNA binding capacity (36) but is also annotated as cold-shock protein CspA, which exerts regulatory effects via RNA binding (37). In addition to RbsR and MsaB/CspA, there are several other transcription factors (MgrA, CcpA, RpiR, SpoVG, CcpE, XdrA, CodY, Rot), two-component regulatory systems (Agr, ArlRS, KdpDE, AirRS, SaeRS) and the alternative Sigma factor B (SigB) known to be involved in regulation of *cap* expression (3, 4). The role of these regulators was mainly deduced from the characterization of single regulatory mutants and in most cases it remains unclear how the regulators affect *cap* expression. In particular, SigB and Agr are believed to act indirectly via other regulatory systems. For instance, the absence of a SigB consensus motif in front of the proposed TSS led to the hypothesis that SigB acts through the SigB-dependent *cap* regulators SpoVG, ArlRS and RbsR (32, 35, 38–40). SigB is a central part of the general stress response, and is activated upon environmental stresses and towards stationary growth phase (41). Its activity is regulated mainly at the post-translational level by a complex regulatory cascade involving RsbW, RsbV and RsbU, encoded within the *rsbUVWsigB* operon.

For the quorum sensing system Agr, it was shown that Agr-mediated *cap* activation occurs via inactivation of the repressor Rot (22). Rot is known to bind to several target genes, however its binding motif is ill-defined (42). Nevertheless, Rot as well as the DNA-binding proteins CodY and SaeR are likely candidates to directly interfere with the P*_cap_* promoter element. CodY represses many metabolic and virulence genes, including the *cap* operon (43, 44), and binding of CodY to the P*_cap_* region has been demonstrated (43, 45, 46). The two component system (TCS) SaeRS regulates a number of virulence factors, with *cap* being one of the few genes repressed (22, 47, 48). With only a poorly conserved SaeR consensus sequence in P*_cap_* it remains unclear whether SaeR exerts its effect on *cap* expression indirectly or via direct promoter interaction.

All-in-all, despite a plethora of work published on *cap* regulation, results are yet inconclusive regarding P*_cap_* architecture and the molecular interference of the different regulatory elements. Here, we re-defined the P*_cap_* promoter structure and re-investigated the main players of *cap* expression. We identified a SigB consensus motif overlapping with the IR structure. In addition, we found a second weak SigA-dependent promoter in the P*_cap_* upstream region, as well as binding sites for three *cap* repressors, CodY, Rot and SaeR, which interfere with SigB-dependent promoter activity. Thus, the early-Off/late-Heterogeneous *cap* expression pattern is a consequence of SigB activity together with repression mediated through the P*_cap_* upstream region. However, negative regulation of CP synthesis in early growth phase is maintained through additional post-transcriptional mechanisms.

## Results

### P_cap_ contains SigA- and SigB-dependent promoters

Previous primer extension analysis revealed a TSS −17 bp upstream of the ATG start codon of *capA* (12). However, RNA-Seq data indicate an alternative TSS further upstream, at position −41 from the *capA* coding region (data not shown), consistent with recent whole genome analyses of TSSs in *S. aureus* (49, 50). To solve this ambiguity, we employed 5’ Rapid Amplification of cDNA Endings (5’ RACE) (Figure 1A). Most of the clones (6/10) revealed a TSS at position −41 bp from the *capA* start codon, which is preceded by a putative SigB consensus sequence. This SigB promoter consists of a conserved SigB −35 motif (GTTTAA) and a −10 region harboring three mismatches (<u>AT</u>GTA<u>A</u> versus GGGTAT) (51). Remarkably, the SigB −35 consensus sequence is located within the IR that is crucial for P*_cap_* activity (12). The identified TSS confirmed our RNA-Seq data and the TSS prediction of Prados et al. (49). In addition, 5’ RACE revealed one clone with a putative TSS at position −128 bp upstream of the *capA* start codon, which was also predicted by Prados et al. (49). A conserved SigA consensus sequence was identified in front of this TSS containing canonical −35 and −10 regions. Interestingly, the TSS −17 bp upstream of the *capA* start codon proposed by Ouyang et al. (12) was not detected. Though one 5’ RACE clone suggested a potential TSS in close proximity, at position −21, this was not preceded by a sigma factor consensus sequence, challenging the presence of a functional promoter. Using 5’ RACE and sequence analysis, we provide evidence for a predominant SigB-dependent promoter and an additional SigA-dependent promoter further upstream in the P*_cap_* promoter element.

**Figure 1:**
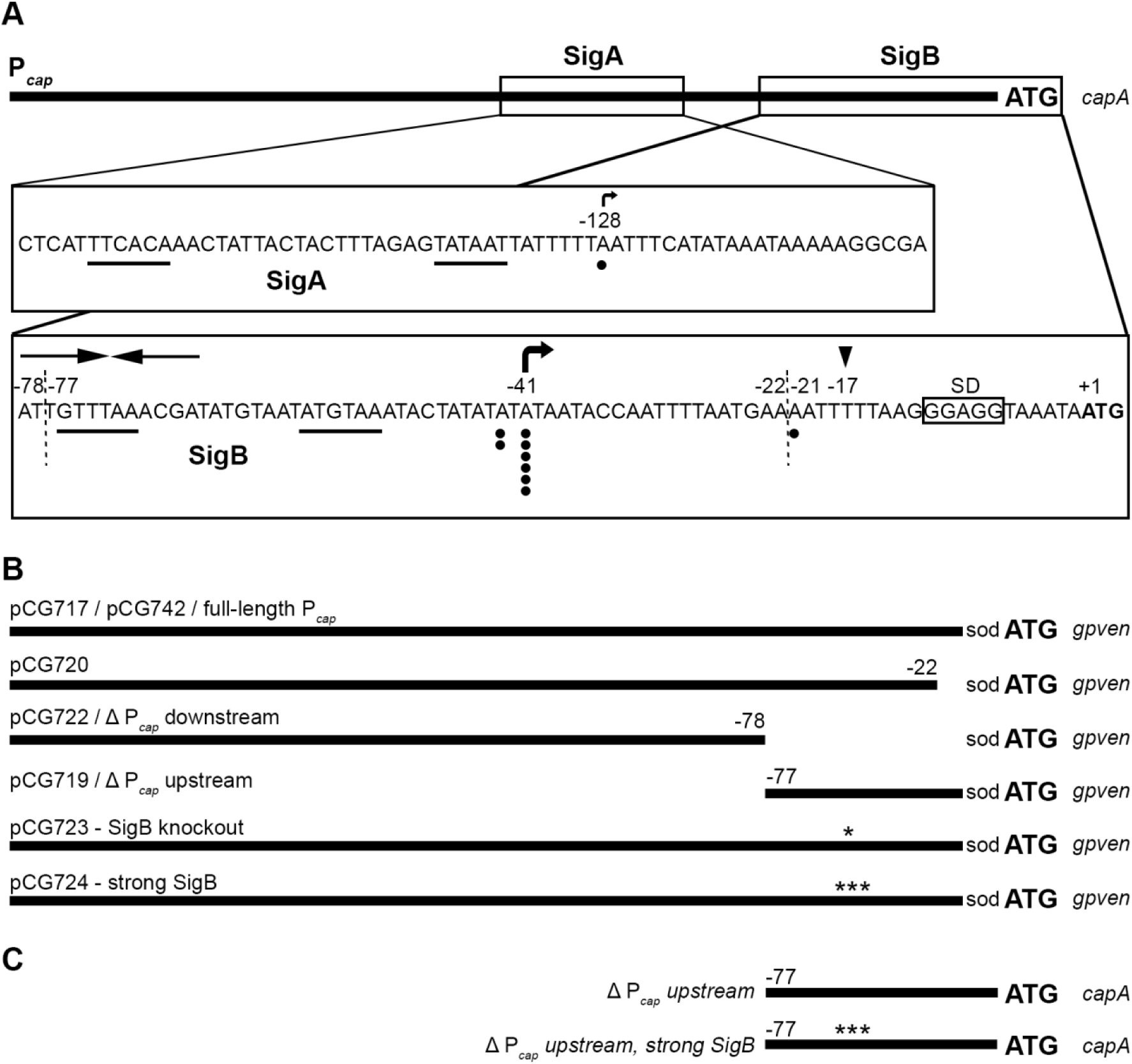
P*_cap_* promoter architecture and P*_cap_* variants employed in this study. (A) P*_cap_* in front of *capA* with magnified SigA- and SigB-promoters. Black dots represent the putative transcriptional start sites (TSSs) suggested by 10 analyzed 5’ RACE clones. Sigma factor −35 and −10 motifs are underlined, bent arrows indicate the corresponding TSSs. The TSS and inverted repeat (IR) structure proposed by Ouyang et al. (12) are marked as black triangle and opposing arrows, respectively. Vertical dashed lines indicate sites of promoter truncation. Numbers mark positions with reference to the ATG of *capA.* The native Shine Dalgano (SD) sequence is labelled and indicated by a box. (B) P*_cap_* promoter fusion variants in front of an artificial ribosomal binding site (sod) and *gpven* gene. Numbers show truncation sites and asterisks indicate point mutations (* −56:G→T; *** −58:A→G, −57:T→G, −52:A→T). (C) Genomic P*_cap_* variants in front of *capA.* Numbers show truncation sites and asterisks indicate point mutations *** −58:A→G, −57:T→G, −52:A→T).

### cap expression is mainly driven by direct SigB regulation

To analyze promoter activities we constructed various P*_cap_*-*gpven* promoter fusions (Figure 1B), including deletions and variations of the putative SigA- and SigB-dependent promoters. Cloning the full-length P*_cap_* in front of *gpven* and an artificial ribosomal binding site (52) (pCG717) resulted in detectable *gpven* expression and was used as reference for all further experiments (Figure 2A). Deletion of the downstream element containing the putative TSSs at position −17 (12) and −21 in pCG720 did not influence *gpven* expression. This supports that there is no active promoter located in this region and that these putative TSSs may have been derived from RNA processing. A construct containing only the SigA-dependent promoter (pCG722) resulted in a low but detectable fluorescence signal, suggesting weak promoter activity. In contrast, a construct containing only the SigB-dependent promoter (pCG719) resulted in strong promoter activity. This indicates dual promoter activity driving P*_cap_* expression: a weak SigA-dependent promoter located in the upstream region plus a main SigB-dependent promoter further downstream. The deletion of the upstream region containing the SigA consensus sequence (pCG719) resulted in promoter activity higher than that of the full-length construct (pCG717). Taken together, these results suggest that despite containing a functional SigA-dependent promoter, the upstream region mainly functions as a repressive element.

**Figure 2:**
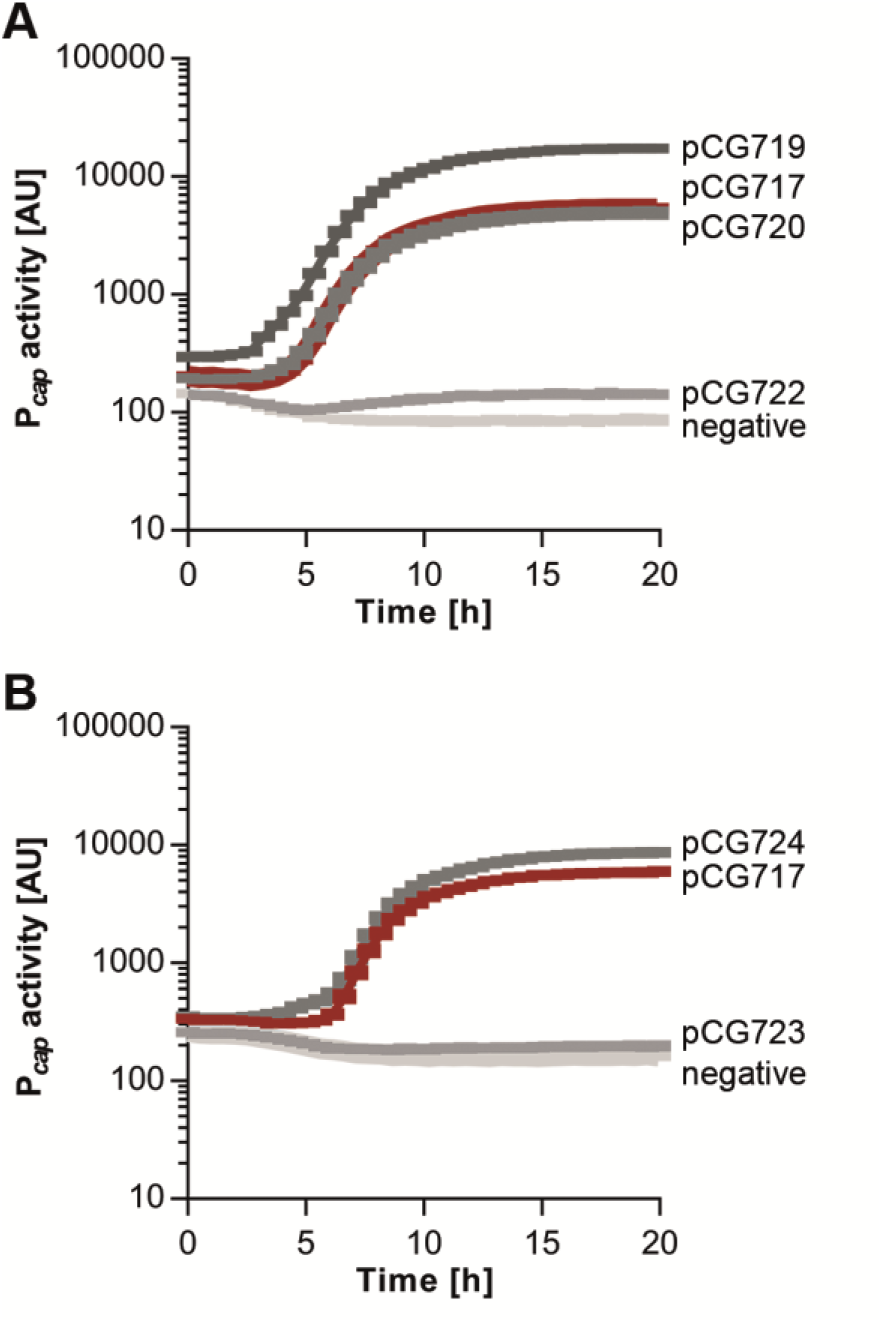
SigB is the main driver of P*_cap_* activity. Promoter activities of different truncated (A) or mutated (B) P*_cap_* fusions in Newman wild-type. The different promoter fusion constructs are described in Figure 1B. The promoter activity of the full-length P*_cap_* promoter fusion (pCG717) serves as reference and is marked in red. Mean gpVenus intensity plus standard deviation of biological triplicates is shown over time.

To further prove that SigB is directly involved in *cap* activation, the expression of the full-length P*_cap_* promoter fusion was measured in wild-type and a *sigB* mutant over time. In addition, constructs where the SigB −10 consensus sequence was mutated to either abolish (pCG723) or enhance (pCG724) SigB affinity were included (Figure 2B). Deletion of *sigB* (data not shown) or a loss-of-function mutation in the SigB consensus sequence eliminated P*_cap_* activity. In contrast, the strong SigB consensus motif further enhanced promoter activity in comparison to the native promoter. These results reveal that *cap* expression is mainly and directly driven by SigB activity.

### Msa activates cap expression by modulating SigB activity

Interestingly, the crucial IR structure that in fact constitutes the SigB −35 motif was shown to be the binding site for two *cap* activators, RbsR and MsaB (35, 36). To elucidate whether and how these regulators interfere with SigB-dependent promoter activity, full-length P*_cap_* promoter fusions were introduced into *msa* and *rbsR* knockout mutants. Under our growth conditions, deletion of *rbsR* showed no effect on P*_cap_* activity (Figure S1).

In the *msa* mutant, P*_cap_* promoter activity was lower than in the wild-type, supporting the finding that MsaB/CspA contributes to *cap* activation (Figure 3A). We hypothesized that MsaB/CspA exerted its effect by modulating SigB activity. To test this hypothesis, we used the double promoter fusion construct pCG742 to simultaneously measure P*_cap_* and P*_asp23_* activity. The P*_asp23_* promoter is widely used as marker for SigB activity (51, 53, 54) and was cloned in front of *gpcer* (52). We found P*_cap_* and P*_asp23_* activity to be highly correlated, in line with the assumption that both are controlled directly by SigB. The activity of both promoters was lower in a *msa* mutant (Figures 3A and B). It was previously shown that MsaB/CspA can bind to *rsbVWsigB* transcript, likely leading to its stabilisation (37). In this case, expression of *sigB* alone should lead to MsaB/CspA-independent regulation. Therefore, we expressed *sigB* from a constitutive promoter in a *rsbUVWsigB* mutant (const. *sigB*). By this means, we also circumvented post-transcriptional regulation of SigB by the RsbUVW phosphorelay. As expected, neither P*_asp23_* activity (Figure 3B) nor P*_cap_* activity (Figure 3A) were affected by *msa* deletion in this background. To exclude additional regulation by direct binding of MsaB/CspA to P*_cap_*, we performed electrophoretic mobility shift assays (EMSAs) with purified MsaB/CspA protein. Even using high amounts of MsaB/CspA protein no band shift was observed (Figure S2). Thus, MsaB/CspA promotes *cap* expression via modulation of SigB activity and not by direct interaction with P*_cap_*.

**Figure 3:**
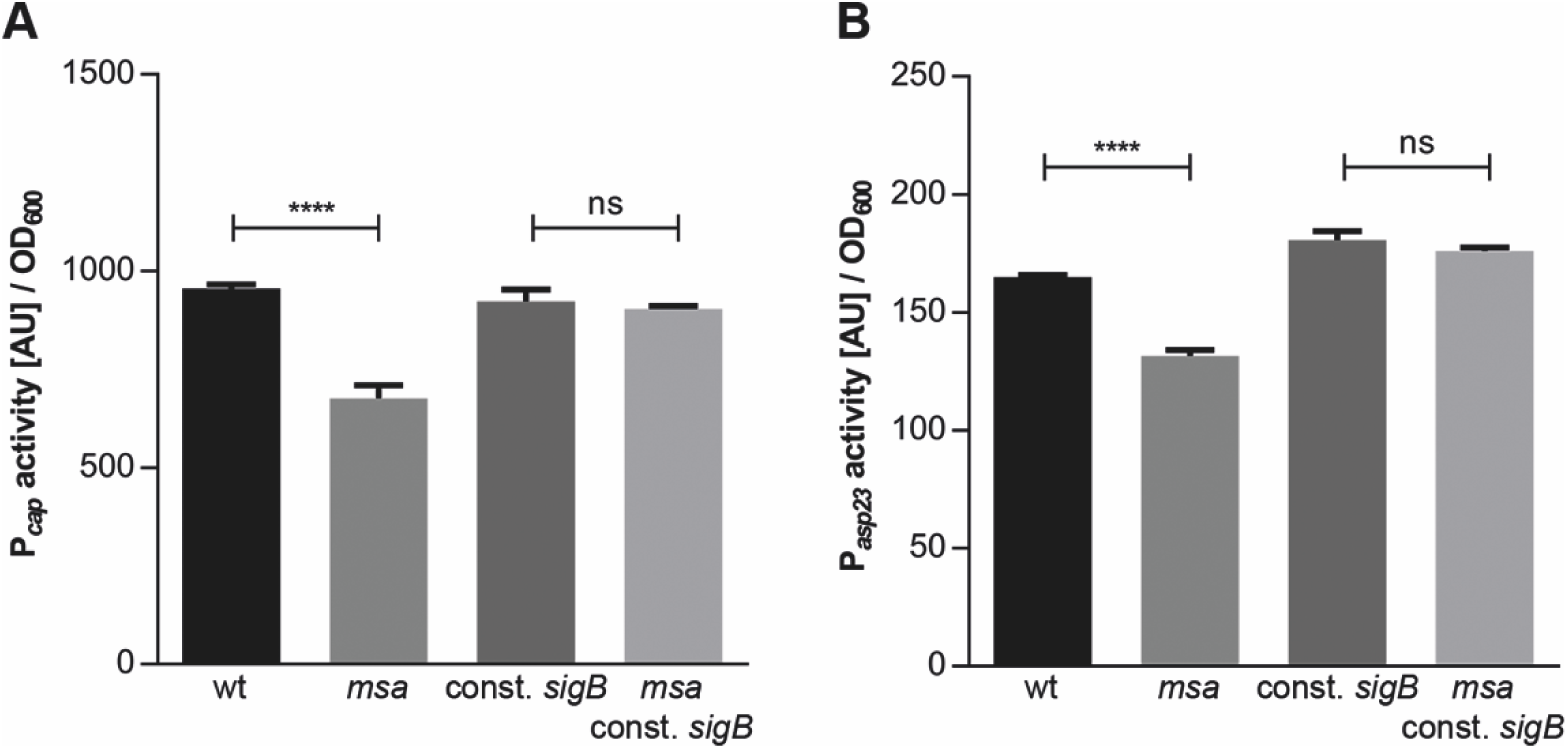
MsaB/CspA activates *cap* expression by modulating SigB activity. Promoter activity of full-length P*_cap_*-*gpven* (A) and P*_asp23_-gpcer* (B) double promoter fusion (pCG742, see Figure 1B) in Newman wild-type and a *msa* mutant with or without constitutive *sigB* expression (const. *sigB*). gpCerulean (P*_asp23_*) and gpVenus (P*_cap_*) intensities are given per OD_600_ after 16 h of growth. Experiments were performed in biological triplicates, error bars represent the standard deviation. Statistical significance was obtained by ordinary one-way ANOVA followed by Tukey’s multiple comparison test (ns: not significant, ****: p<0.0001).

### Upstream promoter region leads to P_cap_ repression via CodY, Rot and Sae

To follow up on the observation that the P*_cap_* upstream region has a repressive function (Figure 2A), we investigated the role of the known *cap* repressors CodY, Rot and Sae. Full-length (pCG717) and truncated (pCG719 and pCG722) P*_cap_* promoter fusions were introduced in *codY*-, *rot*- or *sae*-negative background. Mutation of any of the three regulators resulted in a significant increase of P*_cap_* activity compared with the full-length construct (Figure 4A). The effect of the individual regulators is additive, since in the *sae codY rot* triple mutant the promoter activity is further enhanced compared to the single mutants. If these repressors target the P*_cap_* upstream region, their deletion should not affect promoter activity of a construct lacking this part of the promoter. Indeed, repressor mutations have no or only minor effects on promoter activity of construct pCG719 (Figure 4B). Interestingly, promoter activity of the full-length construct in the triple mutant remained significantly below the level of the upstream truncated construct (Figures 4A and B). This indicates that besides CodY, Sae and Rot, additional repressive factors are acting on the P*_cap_* upstream region.

**Figure 4:**
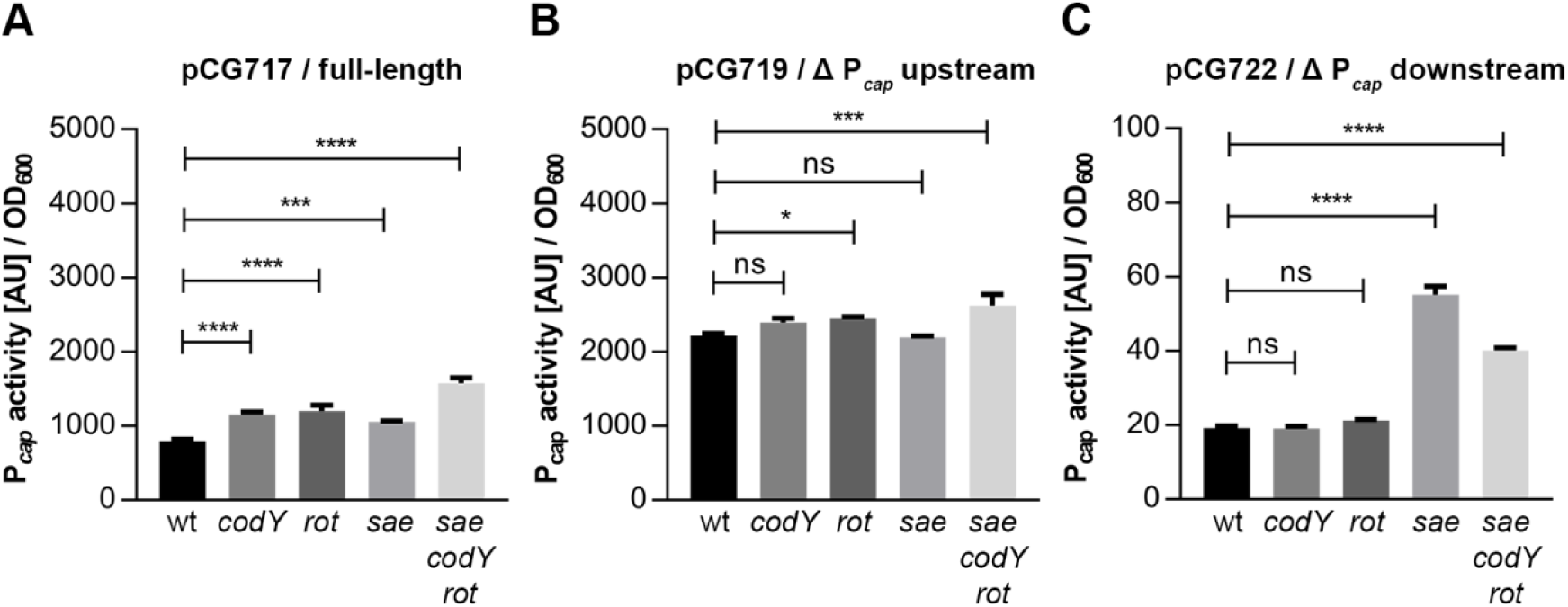
CodY, Rot and Sae repress *cap* expression by interfering with the P*_cap_* upstream region. Promoter activity of different P*_cap_* truncations in Newman wild-type, *codY*, *rot*, *sae* single and triple mutants (A-C). The different promoter fusion constructs are described in Figure 1B. gpVenus intensity is given per OD_600_ after 16 h of growth. Experiments were performed in biological triplicates, error bars represent the standard deviation. Statistical significance was determined by ordinary one-way ANOVA followed by Dunnett’s multiple comparison test (ns: not significant, *: p<0.1, ***: p<0.001, ****: p<0.0001).

We further analyzed whether the repressors also affect the activity of the weak SigA-dependent promoter located in this region (Figure 4C). Remarkably, neither Rot nor CodY showed any influence on promoter activity in a construct only containing the upstream part of the promoter (pCG722). However, mutation of *sae* resulted in increased SigA-dependent promoter activity, even though it remained weak in comparison to the full-length promoter (pCG717). Hence, while all three repressors target the upstream region and affect SigB-dependent promoter activity, only Sae additionally represses the weak SigA-dependent promoter.

### Sae, Rot and CodY repress P_cap_ by binding to the upstream promoter region

We have demonstrated that Sae, Rot and CodY repress *P_cap_*, but it remains unclear whether repression occurs through direct DNA-protein interaction or rather indirectly through the complex regulatory network. To elucidate the nature of the repressors, we performed EMSAs with purified SaeR, Rot and CodY proteins. As SaeR only binds DNA in its phosphorylated state (55), we created a phosphomimetic SaeR with a D51E substitution. Incubation of increasing amounts of SaeR^D51E^ with fluorescence-labelled P*_cap_* upstream fragment (−78 to −344 from *capA*, see Figure 1A) resulted in a retarded protein-DNA complex (Figure 5A), which was not observed using the unphosphorylated native SaeR (Figure 5B). Binding is consistent with a putative SaeR binding motif located between −79 bp and −94 bp upstream of the *capA* start codon (56). Specific binding to the P*_cap_* upstream region was also found for Rot and CodY (Figure 5C and D). These findings are in line with the promoter activities described above showing that Sae, Rot and CodY target the P*_cap_* upstream region (Figure 4A and B). Binding of the repressors to the downstream fragment (+10 to −77 from *capA*, see Figure 1A) is unspecific as band shifts are not eliminated by specific unlabeled competitor (Figure S3 A-D). However, it was previously found that CodY can interact with a region further downstream (+160 to −152 from *capA*, see Figure 1A) reaching into the *capA* coding sequence (45). This region was so far not included in our promoter activity assays or EMSAs. We could confirm CodY binding to this region, when using a 3’ extended downstream fragment (+160 to −77 from *capA*, see Figure 1A) covering the *capA* coding region (Figure S3 E). Together, these results show that SaeR and Rot directly bind to the P*_cap_* upstream region, whereas CodY can bind to two distinct sites, one within the P*_cap_* upstream region, and one further downstream reaching into the *capA* coding region.

**Figure 5:**
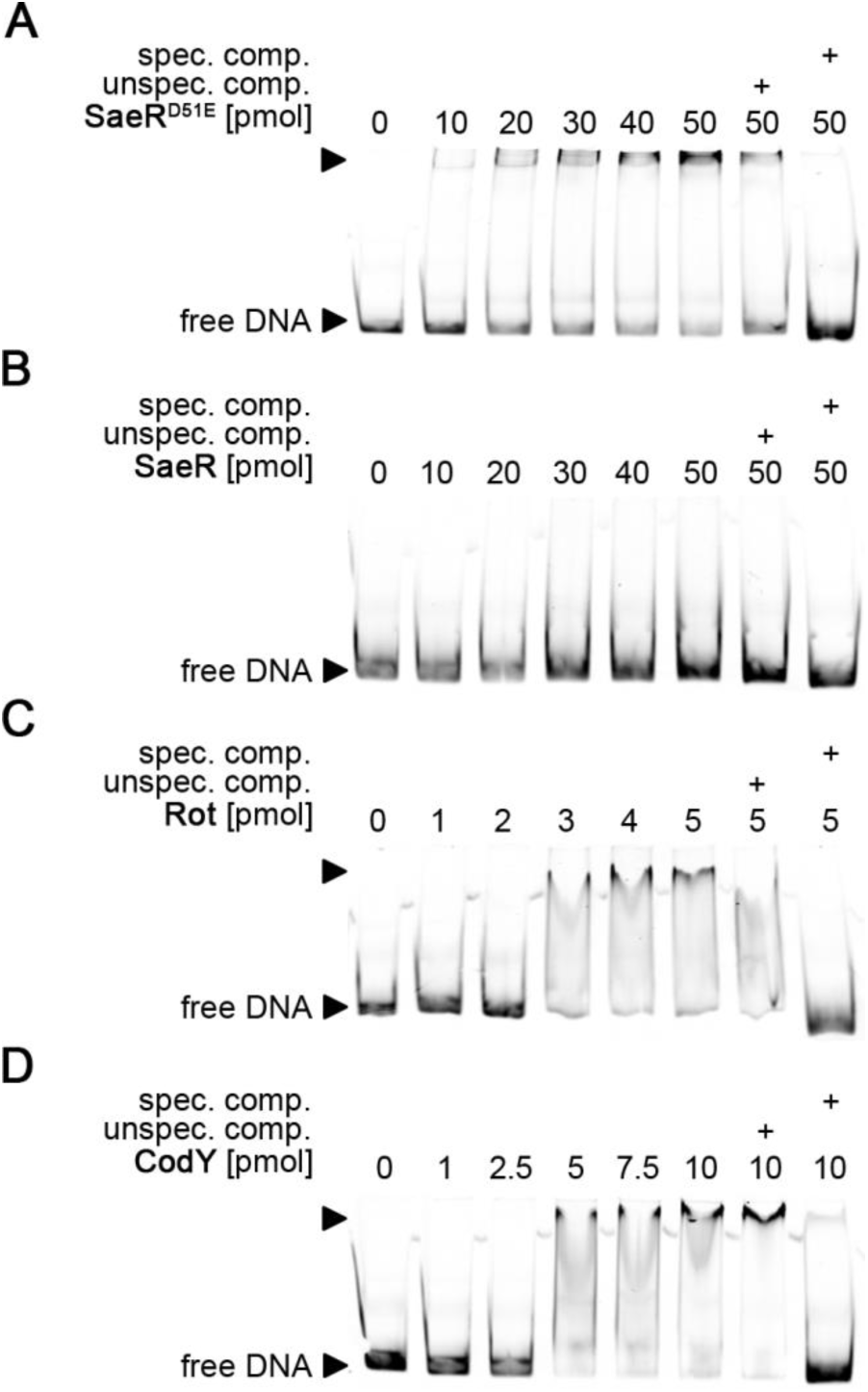
CodY, Rot and SaeR bind directly to P*_cap_* upstream region. Electrophoretic mobility shift assays of purified SaeRD51E (A), SaeR (B), Rot (C) and CodY (D). Increasing amounts of protein were incubated with fluorescently labelled P*_cap_* upstream region (−78 to −344 from *capA*, see Figure 1A). As a control for specificity, shifts were subject to competition with the promoter of the 16 S rRNA gene (unspec. comp.) or unlabelled probe (spec. comp.) in 100 fold excess. Representative pictures from at least three independently performed experiments are shown.

### SigB-dependent regulation and various repressors targeting the upstream region contribute to temporal and heterogeneous CP synthesis

So far we analyzed P*_cap_* promoter activity using artificial promoter fusion constructs. To confirm that our findings translate into CP production we used immunofluorescence (IF) for CP detection. This also allows monitoring the onset of CP production and CP heterogeneity on the single cell level. Cultures were diluted thrice to ensure that all bacteria are actively dividing after inoculation and do not carry residual CP from stationary phase. Bacteria were analyzed throughout growth at different time-points (T_0_ - T_4_). On exponentially-growing bacteria no CP was detectable, but upon reaching stationary phase approximately 30 % of the population became CP-positive, which is consistent with previous results (22). With SigB being a regulator of late genes, we first investigated the effect of constitutive *sigB* expression on CP synthesis during growth. Constitutive *sigB* expression in a *rsbUVWsigB* negative background (const. *sigB*) resulted in earlier onset of CP production and more CP-positive bacteria in stationary phase (Figure 6). In addition, we generated a P*_cap_* mutant in which the upstream promoter region containing the repressor binding sites was chromosomally deleted (*ΔP_cap_ upstream*, Figure 1C). Also in this strain CP production started earlier and more CP-positive cells could be detected towards late growth phases. Of note, the effect of the P*_cap_* upstream region deletion was more profound than that of constitutive *sigB* expression. Furthermore, IF revealed that in bacteria from stationary phase, much of the heterogeneity is reduced in the Δ*P_cap_ upstream* strain and omitted in combination with constitutive *sigB* expression. Thus, it seems to be the combination of repression via transcriptional regulators and SigB-dependent promoter activity that is responsible for the heterogeneous CP expression pattern in stationary phase.

**Figure 6:**
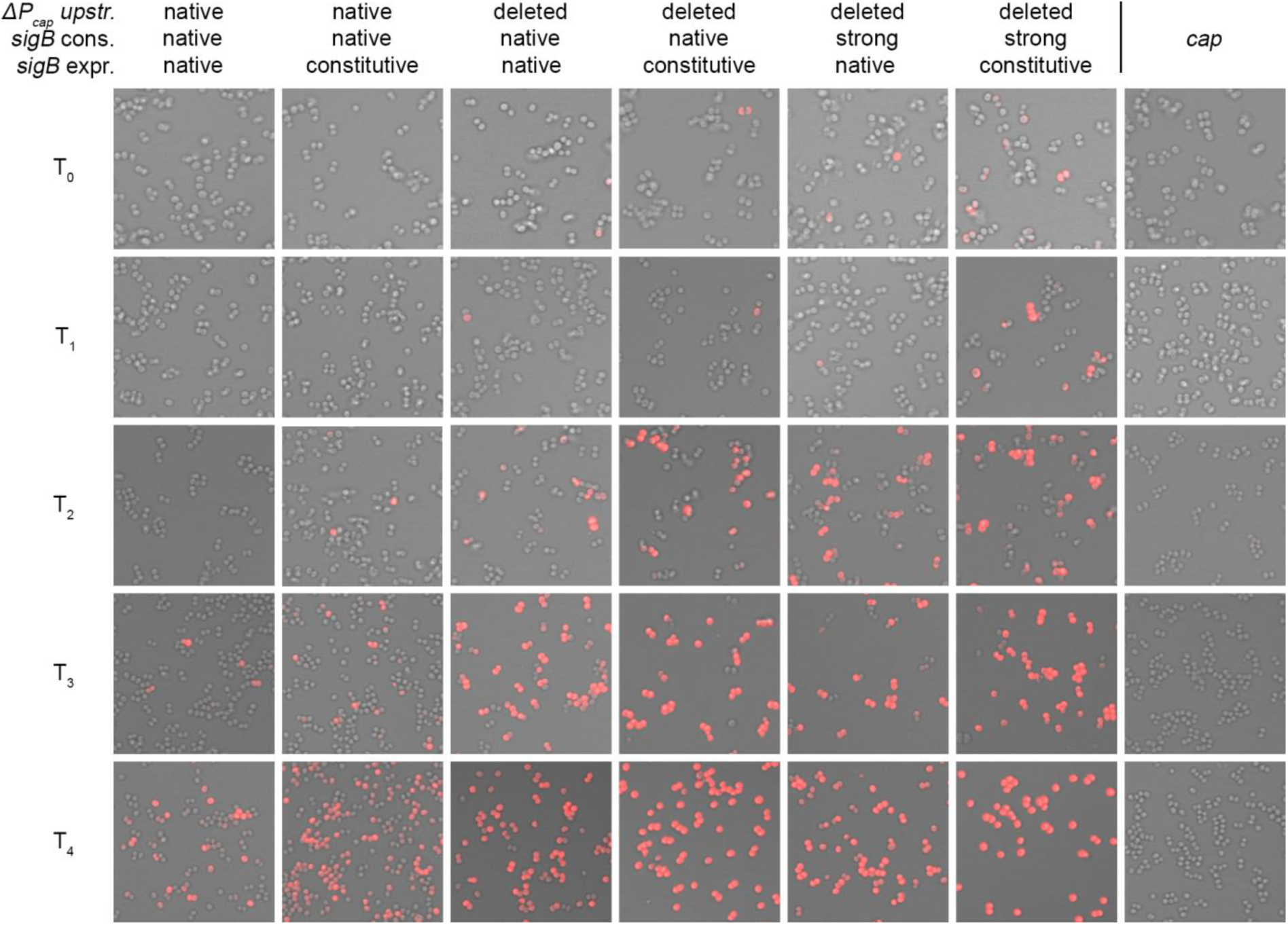
SigB regulation and upstream repressors determine temporal and heterogeneous CP synthesis. Detection of CP production by bacteria from different growth phases via immunofluorescence. The wild-type strain Newman was mutated in order to delete the upstream region (*ΔP_cap_ upstr.*), to contain a strong SigB −10 consensus motif (*sigB* cons.) and/or constitutive *sigB* expression (*sigB* expr.) and grown to defined growth phase T_0_ - T_4._ The different genomic P*_cap_* variants are described in Figure 1C. Representative pictures from at least three independent cultures are shown.

However, throughout all experiments CP expression remained growth phase-dependent. We thought this could be due to the weak SigB promoter of P*_cap_*. Therefore, we additionally altered the SigB −10 region to the conserved SigB −10 motif on the chromosome in the P*_cap_* upstream truncated strain *(ΔP_cap_ upstream, strong SigB*, Figure 1C). Together with constitutive *sigB* expression, this shifted the onset of CP even further towards early growth phase. However, the majority of the bacterial population still remained CP-negative in early logarithmic growth phase.

### CP synthesis is not always correlated to capA transcription

To analyze whether the residual growth phase dependency was correlated to *capA* transcript levels, *capA* mRNA was quantified by qRT-PCR at T_0_ – T_2_ (Figure 7). As expected, in the wild-type strain *capA* expression was strongly repressed in early growth phase. In contrast, constitutive *cap* A expression was achieved in a strain with upstream-deleted P*_cap_* promoter, strong SigB consensus sequence and constitutive *sigB* expression (*ΔP_cap_ upstream, strong SigB*, const. *sigB)*, exceeding that of the wild-type. Of note, CP production is still growth phase-dependent under these conditions (Figure 6), suggesting the existence of post-transcriptional mechanisms regulating CP synthesis.

**Figure 7:**
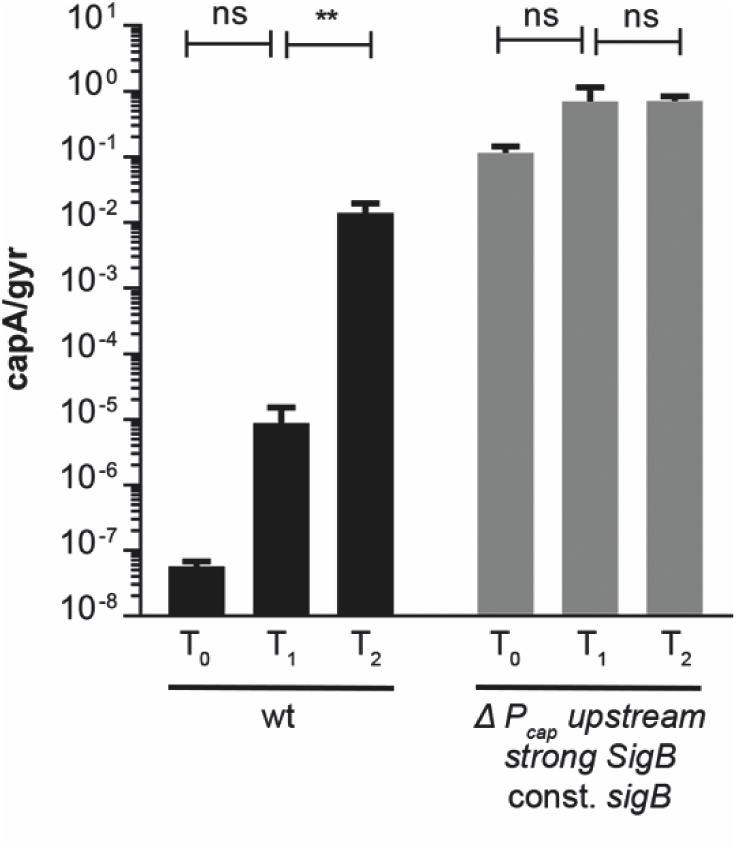
*capA* transcription can be rendered constitutive. Bacterial cells were harvested at different growth phases (T_0_-T_2_) and total RNA was isolated. *capA* transcripts in Newman wild-type and a P*_cap_* upstream truncated strain with strong SigB −10 consensus sequence and additional constitutive *sigB* expression (Newman *ΔP_cap_ upstream, strong SigB*, constitutive *sigB*) were measured by qRT-PCR and normalized to *gyrB.* The genomic P*_cap_* variant is described in Figure 1C. Experiments were performed in biological triplicates, error bars represent the standard deviation. Statistical significance was determined by ordinary one-way ANOVA followed by Tukey’s multiple comparison test (ns: not significant, **: p<0.01).

### P_cap_ regulation is conserved in different S. aureus strains

So far all experiments have been performed in strain Newman which is special due to a hyperactive SaeRS system (57). To validate our findings in a different *S. aureus* background we chose the widely studied community-acquired Methicillin-resistant *S. aureus* strain USA300 JE2. This strain has an acapsular phenotype due to three crucial conserved mutations in the P*_cap_* promoter region and in the coding regions of *cap5D* and *cap5E* (15). Therefore, we first generated derivatives in which we either only repaired the mutation in P*_cap_* (*P_cap_ repaired*) or all three mutations (*cap repaired*) (Figure 8). In line with previous observations, the USA300 JE2 wild-type shows an acapsular phenotype and the repair of the mutation in P*_cap_* alone is not sufficient to enable CP production (15). Only when all three mutations were repaired, USA300 JE2 was capable of producing CP, following the same peculiar expression pattern as strain Newman: CP-positive cells were observed towards late growth phase and CP expression was highly heterogeneous. Upon deletion of the P*_cap_* upstream region and introduction of the fully conserved SigB consensus sequence (*cap repaired*, Δ*P_cap_ upstream, strong SigB*) CP-positive cells were detected earlier and in late growth phase all cells were CP-positive. Therefore, the CP expression pattern of USA300 *cap repair* and its regulation closely resembles that of Newman.

**Figure 8:**
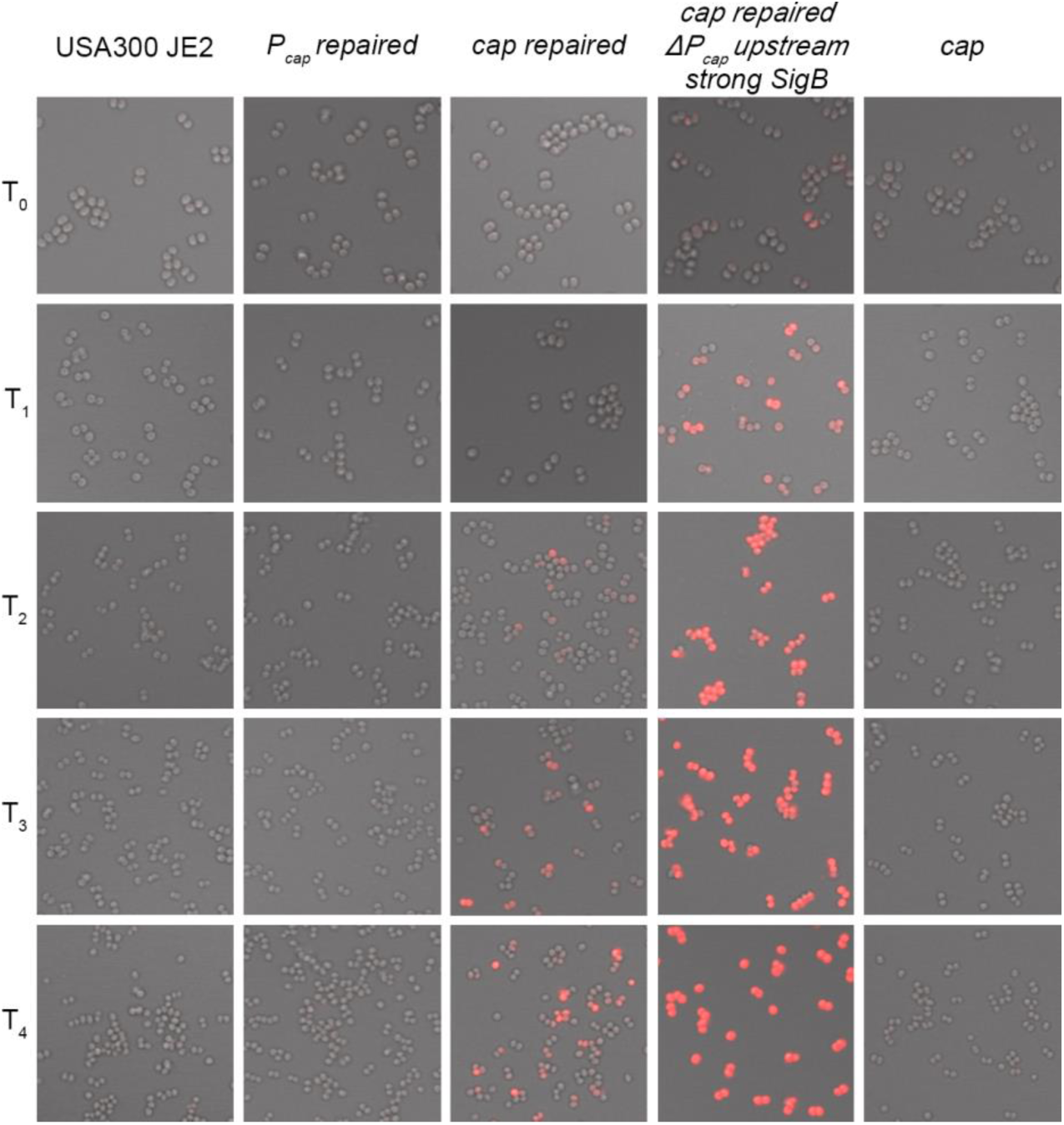
CP production in strain USA300 JE2. Detection of CP in bacteria from different growth phases. USA300 JE2 wild-type and mutants in which either the mutation in P*_cap_* alone (*P_cap_ repaired*) or in the whole *cap* locus (P*_cap_*, *capD, capE; cap repaired*) are repaired were grown to defined growth phase T_0_ - T_4_ and CP was detected by immunofluorescence. In addition, the effect of P*_cap_* upstream truncation and a strong SigB −10 consensus sequence (*cap repaired, ΔP_cap_ upstream, strong SigB*) was analyzed. This genomic P*_cap_* variant is described in Figure 1C. Representative pictures of at least three independent cultures are shown.

## Discussion

CP protects *S. aureus* against phagocytosis, but also hampers adherence to endothelial cells and matrix proteins. It is believed that heterogeneity of CP expression has evolved to provide better adaptability of the bacterial population during infection and colonization (22). Apart from this heterogeneous phenotype, CP production is strongly growth phase-dependent, with encapsulated cells found only towards stationary growth phase (22). During the last decades many regulatory proteins have been shown to have an impact on CP production, forming a complex regulatory network. However, to date it is not known which regulator is mainly responsible for this peculiar expression pattern. Here, we revisited the role of important molecular determinants of P*_cap_* activity, summarized in Figure 9.

**Figure 9:**
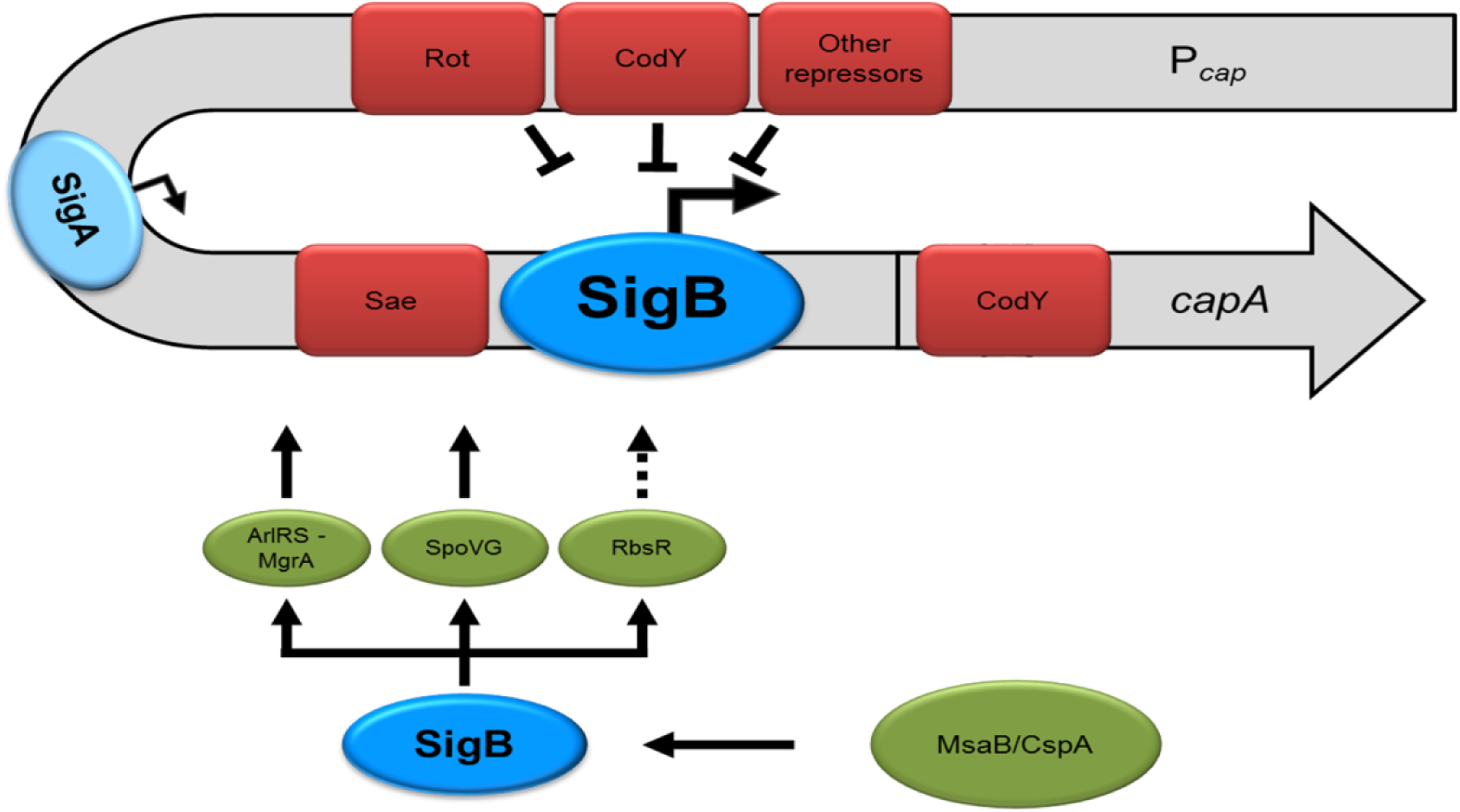
Transcriptional regulation of *cap* expression. *Cap* expression is mainly driven from a SigB-dependent promoter. However, there is an additional weak SigA-dependent promoter further upstream. In addition, the P*_cap_* upstream region is targeted by the *cap* repressors SaeR, CodY, Rot and others. Of note, Rot and CodY only interfere with SigB-dependent promoter activity, whereas Sae is able to repress both promoters. A predicted SaeR binding motif is located between the SigA- and SigB-dependent promoters. P*_cap_* contains a second CodY binding site reaching into the coding region of *capA*. SigB activity is modulated by MsaB/CspA, promoting *sigB* transcript stability. Consequently, MsaB/CspA likely also affects indirect SigB-dependent P*_cap_* activation through ArlRS-MgrA, SpoVG and RbsR. However, for RbsR we were unable to prove an activating effect on *cap* expression.

### cap expression is mainly driven by SigB activity

We show that the main P*_cap_* promoter is SigB-dependent. This is based on i.) 5’ RACE results revealing a new TSS downstream of a conserved SigB consensus sequence with minor mismatches in the −10 region; ii.) only very weak promoter activity in a *sigB* mutant or upon mutation of the −10 consensus sequence; iii.) higher P*_cap_* activity when the SigB −10 region matches the proposed SigB consensus sequence; iv.) a strong correlation of P*_asp23_* and P*_cap_* promoter activity. Of note, the SigB −35 region is directly located within the IR region shown to be crucial for *cap* expression (12). Thus, the requirement of this structure can be simply explained by its function as a SigB binding site. A positive effect of SigB on *cap* expression was previously shown. However, it was believed to be mediated via the SigB-dependent c*ap* activators SpoVG, ArlRS-MgrA and RsbR (32, 35, 38–40). Our results highlight the importance of SigB for *cap* expression and clearly demonstrate that its main impact is through direct SigB-dependent promoter activation. Nevertheless, other SigB-dependent *cap* regulators may contribute to the fine tuning of *cap* expression and amplify SigB dependency.

### MsaB/CspA activates cap expression by modulating SigB activity

It was previously shown that the SigB −35 consensus sequence functions as binding site for two *cap* activators RbsR and MsaB (35, 36). We did not observe an effect of RbsR on P*_cap_* activity, neither in wild-type nor in a *sae* negative background. RbsR likely functions as metabolic sensor, and thus the discrepancy with the results of Lei and Lee (35) could be due to differences in growth conditions.

We reproduced the activating effect of MsaB/CspA on *cap* expression, but demonstrated that it is mediated by modulation of SigB activity: *msa* deletion negatively affected P*_cap_* and P*_asp23_* activity and constitutive expression of *sigB* rendered both promoters insensitive to MsaB/CspA. Moreover, in contrast to Batte et al. (36), no detectable MsaB/CspA binding to P*_cap_* was observed, challenging the role of MsaB/CspA as a classical transcription factor. Strong interference of MsaB/CspA with SigB activity is consistent with previous findings that MsaB/CspA increases expression of *sigB* and its target genes (58, 59). It was recently shown that MsaB/CspA binds *rsbVWsigB* mRNA, thereby increasing transcript stability (37). With respect to direct P*_cap_* regulation, our results indicate that under the conditions employed, SigB is the only protein binding to the IR/SigB −35 consensus sequence.

### Cap expression is modified by upstream SigA promoter and repressor binding sites

Next to the main SigB-dependent promoter we identified by 5’ RACE an additional weak SigA-dependent promoter with conserved SigA −35 and −10 consensus sequences. However, the SigA-dependent promoter seems to play a minor role in *cap* expression, with its activity being mainly detectable upon deletion of *sae*. The reason for such low promoter activity is unclear and might be due to sub-optimal structure and spacing of the SigA consensus in P*_cap_*. (60, 61). Furthermore, the average activity of SigA promoters has been found to be generally lower than that of SigB promoters (50). For any given promoter sequence, changes in temperature, salt and solute concentrations, as well as protein factors and ligands can affect its kinetics by 10 - 1000 fold or more (61). Therefore, we cannot rule out that under certain conditions the SigA-dependent promoter gets activated. This may be the case during infections with strains from the USA300 lineage. Amongst others, USA300 strains carry a point mutation in P*_cap_* located within the SigB consensus motif defined here. Interestingly, strains harboring only this mutation seem to produce CP under infectious conditions (16), which might be attributed to activation of the weak SigA promoter.

Besides containing a SigA-dependent promoter, we observed that the P*_cap_* upstream region is targeted by many transcriptional factors. For the *cap* repressors SaeR, CodY and Rot, direct interaction with the P*_cap_* upstream region was demonstrated by EMSAs. *cap* repression was confirmed by promoter activity assays, suggesting that the main function of the upstream promoter region is to reduce SigB-dependent promoter activity. While Sae, CodY and Rot all interfered with SigB-dependent promoter activity, only Sae additionally repressed activity of the SigA-dependent promoter. This is in line with a predicted SaeR binding site located between the two promoters (56). We hypothesize that the repression of SigB-dependent promoter activity occurs via steric interference, whereas the SigA-dependent promoter activity might be repressed via a roadblock mechanism. Nonetheless, the molecular mechanism for the long-distance effect of Rot and CodY on the SigB-dependent promoter activity remains to be elucidated. One may speculate that secondary structures of the promoter bring these two regulators in close proximity to the SigB consensus motif, allowing them to interfere with SigB binding. It is well known that DNA structural elements like supercoiling are involved in the control of bacterial gene expression (62), and *cap* expression was indeed shown to be supercoiling sensitive (63).

Of note, there are two CodY binding sites within the *cap* locus. One reaching into the coding region of *capA*, consistent with previous findings (45), and one in the P*_cap_* upstream region. Functional assays using truncated promoter fusion constructs indicate that the upstream binding site alone is sufficient for *cap* repression.

### SigB activity and upstream repressors determine temporal/heterogeneous CP synthesis

We showed that SigB-dependent regulation and the regulators targeting the upstream region contribute to the temporal pattern of CP production and to its phenotypic heterogeneity. Congruent with SigB being a known activator of late genes (38, 39, 41, 50), we observed earlier onset of CP production and more CP-positive cells in stationary phase upon constitutive *sigB* expression in a *rsbUVWsigB* negative background. However, the greater impact on temporal and heterogeneous CP production was observed upon deletion of the upstream promoter region resulting in single CP-positive cells already in early exponential growth phase and almost all cells being CP-positive in later growth. The effects of constitutive *sigB* expression and P*_cap_* upstream deletion were additive, and only in combination the phenotypic heterogeneity could be completely abolished in stationary growth phase. This suggests that the regulators targeting the upstream region of P*_cap_* are mainly responsible for heterogeneous CP production, with the residual heterogeneity resulting from variable SigB activity within cells. These data support the prediction by Sharon et al. stating that more transcription factor binding sites result in noisier promoters (64). A similar pattern of CP production was shown in the USA300 background, indicating a conserved regulatory mechanism.

Of note, even upon P*_cap_* upstream deletion and constitutive *sigB* expression CP synthesis remains growth phase-dependent which is not reflected at the transcriptional level, indicating that further post-transcriptional levels of regulation are in place. These might be required as CP synthesis is linked to the metabolic status of the cell. For instance, UDP N-acetylglucosamine used for CP biosynthesis is mainly derived from gluconeogenesis, which naturally occurs when glucose becomes limited towards later growth phases (65). In addition, CP, peptidoglycan and wall teichoic acids synthesis make use of the universal bactoprenol carrier lipid, which could become limited in earlier growth phases (66). This coordination of the different cell wall polymers synthesis was recently shown to involve reversible protein phosphorylation of the capsular biosynthesis gene products CapM and CapE (10).

In summary, our results show that CP synthesis is tightly controlled at the transcriptional level. However, post-transcriptional mechanisms are also in place to avoid conflict between precursor usage by the CP synthesis machinery and the synthesis machinery of other cell-wall glycopolymers in growing bacterial cells. Further in-depth studies are required to fully understand this regulation and to increase the potential of CP as prospective target for novel anti-infective strategies.

## Methods

### Bacterial strains and growth conditions

Strains and plasmids are listed in tables S1 and S2. For overnight culture, strains were grown in Luria-Bertani (LB) medium (low salt) with appropriate antibiotics (10 μg mL^−1^ chloramphenicol (Cm10), 10 μg mL^−1^ erythromycin, 50 µg mL^−1^ kanamycin, 3 μg mL^−1^ tetracycline) at 37 °C and 200 rpm. Day cultures were inoculated from overnight cultures to an OD_600_ of 0.05 and were grown without antibiotics at 37 °C and 200 rpm.

### Strain construction

All plasmids and oligonucleotides are listed in tables S2 and S3, respectively. The transposon mutants Newman *rbsR* and Newman *sae rbsR* were constructed by phage transduction of the transposon insertions from the Nebraska transposon mutant NE425 to Newman and Newman-29 and then verified by PCR.

Strains Newman *msa*, Newman *ΔP_cap_ upstream*, Newman *ΔP_cap_ upstream, strong SigB*, USA300 *P_cap_ repaired*, USA300 *cap repaired* and USA300 *ΔP_cap_ upstream, strong SigB* were created using the temperature-sensitive pIMAY plasmid for allelic exchange (67). Corresponding homologous regions were PCR-amplified as indicated in table S3 and inserted into the *Eco*RI-linearized pIMAY backbone via Gibson assembly. The resulting plasmids were verified by PCR using primers piMAYcontrolfor and piMAYcontrolrev and sequencing, and then electroporated into strain RN4220 and transduced into the target strains. Allelic exchange was performed as described before (67) with few alterations. Briefly, a single colony was homogenized in 200 µL TSB and 50 µL of serial dilutions (10^−1^ – 10^−3^) were plated on TSA-Cm10 and incubated overnight at 37 °C. Large colonies were picked and sub-cultured on TSA-Cm10 at 37 °C. Integrants were confirmed via PCR using primers piMAYcontrolfor and piMAYcontrolrev as well as a primer pair flanking the individual homologous regions (Table S3). One integrant colony was used to inoculate an overnight culture in 10 mL TSB grown at 28 °C without chloramphenicol, and later plated on TSA containing 0.7 - 1 µg mL^−1^ Anhydrotetracycline (ATc). After 48 h of growth at 28 °C, colonies were picked on both blood and TSA-Cm10 plates and incubated at 37 °C. Cm10-sensitive colonies were screened by PCR with the oligonucleotides mentioned above and by sequencing to identify clones containing the desired mutation.

The *rsbUVWsigB* mutants were obtained using the temperature-sensitive shuttle vector pBASE6 (68). Replacement was introduced by creating PCR products using RN6390 as a template and primer pairs MazSIG-for and Hybrid-MazSIG-rev, as well as MazSIGrev and Hybrid-MazSIG-for for the homologous regions. The tetracycline resistance cassette was amplified from pCG75 using primers TetMfor and TetMrev. PCR products were linked using primers BglIIMazSIGfor and SalIMazSIGrev. The obtained amplicon was cloned into *Bgl* II and *Sal* I digested pBASE6. The resulting plasmid pCG331 was introduced into the restriction-deficient strain RN4220 and transduced into *S. aureus* target strains. Mutagenesis was performed as described elsewhere (69). The deletion was verified by PCR with oligonucleotides flanking the deleted region and within the resistance cassette.

For constitutive *sigB* expression under the control of the P*_SarA_* promoter, the *sigB* gene including its native ribosomal binding site was amplified from Newman genomic DNA using primers pCG795gibfor and pCG795gibrev and inserted into *Eco* RI and *Asc* I digested pJL78. The resulting plasmid pCG795 was verified via PCR and sequencing prior to electroporation into strain RN9011 and transduction into target strains. Correct insertion into the SaPI-site was verified via PCR using primers SaPIintfor and SaPIintrev.

The *sae codY rot* triple mutant was obtained by transduction of the Δ*codY::tetM* mutation from RN4220-21 (44) into Newman-29, followed by transduction of the *rot::bursa aurealis* transposon insertions from the Nebraska transposon mutant NE386.

For the generation of a *cap* mutant in USA300 JE2, pCG132 was transduced from RN4220 pCG132 (33) into USA300 JE2 using phage Φ11.

### Promoter fusion constructs

Two tandem sequences, each comprising a promoter of interest (P*_gene_*) followed by a strong ribosomal binding site (RBS), genes encoding for fluorescent protein gpVenus (*gpven*) or gpCerulean (*gpcer*) (52) and a terminator (ter) sequence, were designed *in silico*. Restriction sites were designed to flank both promoter replacements. The entire cassette encompassing P*_cap_*-RBS-*gpven*-ter—P*_tagH_*-RBS-*gpcer*-ter was also flanked by restriction sites. The dual promoter-reporter protein fusion cassette was synthesized by Life Technologies GmbH and provided in a pUC18-like *E. coli* vector backbone (pMA-RQ, geneart plasmid). The dual promoter-reporter fusion cassette was sub-cloned into pCG246 (70) using restriction sites *Sph* I and *Nar* I. The resulting plasmid pCG570 was verified by restriction digestion (*Bam*HI) and using the primers pCG246for, pCG246rev, Pcap_rev and PtagHrev for sequencing. The plasmid pCG570 was introduced into RN4220 and then transduced into strain Newman.

Promoter truncations and mutations were generated with the Q5 Site-Directed Mutagenesis Kit from NEB according to the manufacturer’s instructions. All primers used are enlisted in table S3. For plasmids pCG742, pCG815 and pCG816, P*_tagH_* was replaced with P*_asp23_* via Gibson cloning. The *asp23* promoter region was amplified from Newman genomic DNA using primers pCG657gibfor and pCG657gibrev and inserted into *Bam* HI and *Eco* RI digested pCG717, pCG719 andpCG769.

### RNA isolation

A day culture was inoculated from an overnight culture and grown to an OD_600_ of 0.3. The day culture was then used to inoculate a second day culture to an OD_600_ of 0.05, which was grown to an OD_600_ of 0.3 and used again for sub-culturing. When the third day culture reached an OD_600_ of 0.3, 10 mL of cell suspension was harvested (T_0_). After additional 2 h (T_1_) and 4 h (T_2_) of growth, 5 mL of *S. aureus* cells were harvested. Cells were lysed in 1 mL TRIzol reagent (Life Technologies, Germany) with 0.5 mL of zirconia-silica beads (0.1 mm diameter) in a high-speed homogenizer. RNA was isolated as described in the instructions provided by the manufacturer of TRIzol (Life Technologies).

### 5’ Rapid Amplification of cDNA Endings (5’ RACE)

5’ RACE was performed as described previously using strain Newman pCG717 (71). Briefly, isolated RNA from T_2_ was treated with MICROBExpress (Ambion) in order to remove rRNAs. After treatment with Cap-Clip Phosphatase (Biozym) to remove pyrophosphate at the 5’ prime end of the native transcripts, a specific RNA 5’ adapter (Table S3) was ligated to the RNA. After phenol/chloroform extraction and ethanol precipitation, the RNA was subjected to reverse transcription using oligonucleotide YFPCFPpolymorrev. Nested PCR was performed using oligonucleotides Race2 and Racecapnestedrev (Table S3). The PCR amplicon was cloned into pCRII-TOPO (Invitrogen) following the manufacturer’s instructions. Single clones were analyzed via PCR using primers Race2 and Racecapseqrev and the PCR products of 10 clones were sequenced with primer Racecapseqrev.

*qRT-PCR*. qRT-PCR to quantify *cap* and *gyr* mRNA was performed using the QuantiFast SYBR-Green RT-PCR kit (Qiagen). Standard curves were generated using 10-fold serial dilutions (10^4^ to 10^8^ copies) of specific *in vitro* transcribed RNA standard molecules (72). The number of copies of each transcript was determined with the aid of the LightCycler software and *cap* mRNA was expressed in reference to copies of *gyrB*.

### Growth curves and fluorescence measurements

Overnight cultures were diluted to an OD_600_ of 0.05 and 200 µL were loaded onto a 96-well U-bottom plate (Greiner Bio-One). Continuous absorbance and fluorescence were monitored with a TECAN Infinite 200 microplate reader every half an hour with shaking at 37 °C. Absorbance was measured at 600 ± 9 nm, gpVenus was excited at 505 ± 9 nm and emission was measured at 535 ± 20 nm, and gpCerulean was excited at 434 ± 9 nm and emission was measured at 485 ± 20 nm. Optical densities measured in microplate reader are not linear over the extended growth-cycle. To address this problem, we simultaneously measured dilutions of an overnight culture in the plate reader and the Ultrospec™ 2100 pro UV/Vis spectrophotometer (Pharmacia). From the plotted values a conversion formula was calculated and used to correct the optical densities measured in the microplate reader.

### Protein expression

Plasmid pCWsae106 containing SaeR^D51E^ was created by overlapping PCR employing the oligonucleotides saetetfor1 and 1680I29Gluok as well as 1684U31 and saetetfor1. The amplicon was inserted into pCR2.1. pCWsae106 was then transformed into RN4220 and transduced into strain Newman-29. Coagulase assays were performed to verify the function of the phosphomimetic SaeR^D51E^. Strain Newman-29 pCWsae106 was used as a template to generate the insert for the SaeR^D51E^ expression vector with the primers pCG791gibfor and pCG791gibrev. The amplicon was inserted into *Bam*HI linearized pET15b via Gibson assembly. All other expression vectors were generated accordingly using primers and template DNA as indicated in table S3. All vectors were verified via PCR and sequencing using primers pET15bfor and pET15brev.

A 10 mL LB culture containing 100 µg mL^−1^ Ampicillin was inoculated with freshly transformed *E. coli* BL21 DE3 cells and incubated for 6 h at 37 °C and 200 rpm. For the expression culture, 1 L LB medium supplemented with 100 µg mL^−1^ Ampicillin and IPTG or D(+)-lactose-monohydrate (for expression conditions see table S4) was inoculated with 10 mL day culture and incubated at 16 °C and 200 rpm overnight (16 - 18 hours). Cells were harvested (20 min, 2000 × g, 4 °C) and resuspended in 30 mL ice-cold HisTrap binding buffer (20 mM Na_2_HPO_4_, 500 mM NaCl, 40 mM imidazole, pH 7.4, sterile-filtered and degassed) supplemented with 10 μg/mL DNAse (Roth) and cOmplete protease inhibitor cocktail (Roche). Cells were lysed using a French press at 1000 psi. The lysate was centrifuged (236 982 × g, 45 minutes, 4 °C), and the clear supernatant was filtered (0.22-μm pore size) before being loaded onto a 1-mL HisTrap HP column (GE Healthcare Life Sciences) equilibrated with HisTrap binding buffer. Purification was performed with an ÄKTA purification system (GE Healthcare Life Sciences), and elution was carried out with an imidazole gradient to a final concentration of 500 mM.

For SaeR^D51E^ batch purification using Ni-NTA agarose resin (Qiagen) equilibrated with HisTrap binding buffer was performed. After incubation for 1 h at constant over-head rotation at 4 °C the Ni-NTA agarose was centrifuged at 500 × g for 2 min at 4°C and washed thrice with 37.5 mL HisTrap binding buffer. Elution was performed at increasing imidazole concentrations (250/300/350/500 mM).

Column and batch fractions were analyzed by SDS-PAGE, and the fractions containing the protein of interest were collected, concentrated and rebuffered into EMSA buffer (10 mM Tris-HCl, 50 mM KCl, 5 mM MgCl_2_, 10 % glycerol, pH 7.4) using Amicon® ultra centrifugal filters (ULTRACEL® 10K/30K) according to the manufacturer’s recommendations. To determine the protein concentration of purified proteins, a Pierce™ BCA assay (Thermo Fisher) was performed following the manufacturer’s protocol for a microplate procedure. The absorbance was measured with the TECAN Infinite 200 microplate reader. Purified proteins were stored in aliquots at −20 °C.

### Electrophoretic Mobility Shift Assay (EMSA)

The primers used for EMSAs are listed in table S3. DNA probes were PCR amplified from strain Newman with fluorescently labelled primers (DY-781, absorption: 783 nm, emission: 800 nm) and purified with the innuPREP PCRpure Kit from Analytik Jena AG according to the manufacturer’s instructions. In a volume of 20 µL, 30 fmol DNA probe were mixed with various amounts of protein. For competition experiments, unlabelled DNA fragment was used as a specific competitor and a 161 bp DNA fragment containing the promoter of the 16S rRNA gene was used as unspecific competitor. Competitors were obtained via ethanol precipitation after PCR and added in 100 fold excess. After incubation for 20 min at room temperature, samples were analyzed by non-denaturing native 6% TBE polyacrylamide gel electrophoresis at 75 V for 90 minutes in 1x TBE buffer. The fluorescently-labelled DNA probes were visualized in the polyacrylamide gel with the Odyssey® infrared imaging system (LI-COR) and the Image Studio 4.0 software.

### Immunofluorescence (IF) and promoter fusion microscopy

For synchronization, three subsequent day cultures were inoculated to an OD_600_ of 0.05, grown to an OD_600_ of 0.3 and sub-cultured twice. When the third sub-culture reached an OD_600_ of 0.3 (T_0_) and after additional 2 h (T_1_), 4 h (T_2_), 6.5 h (T_3_) and overnight growth (T_4_) cells were harvested by centrifugation at 3800 × g for 10 min at 4 °C. Roughly 1.3 × 10^8^ bacteria were suspended in 1 mL fixation solution (3.7 % formaldehyde in 1x PBS) and incubated with gentle mixing for 15 min at room temperature. Wells of Ibi-treat µ-slide angiogenesis slides (ibidi®) were loaded with 32 µL of cell suspension and centrifuged at 600 × g for 6 min. The cells were washed with 1 × PBS and protein A was blocked by incubation with pre-adsorbed human serum (diluted 1:10 in 1x PBS/0.1 % Tween 20) for 1 h. After blocking, the slides were washed thrice for 5 min each with PBS/Tween 20 followed by incubation with rabbit serum raised against CP5 (1 h, diluted 1:200 in PBS/Tween 20). The slides were washed thrice with PBS/Tween 20 followed by incubation with the secondary antibody Cy3-conjugated F(ab)2 goat-anti-rabbit IgG (Dianova, Hamburg) (diluted 1:500 in PBS/Tween 20, 1 h). For generation of pre-adsorbed human serum and antibody generation, see (22). After washing the cells thrice with PBS/Tween 20 each slide was finally mounted using ibidi® fluorescence mounting medium.

Microscopy images were acquired in the confocal mode of an inverted Zeiss LSM 710 NLO microscope equipped with a spectral detector and a Zeiss Plan-Apochromat 63x/1.40 oil DIC M27 objective and ZEN Black software. The following excitation wavelength, laser sources and detection spectra were used for immunofluorescence: Cy3: Ex: 561 nm/ DPSS laser/ Em: 566 – 702 nm and for promotor activity measurement: gpVenus: Ex: 514 nm/ argon laser/ Em: 519 – 554 nm, gpCerulan: Ex: 405 nm/ diode laser/ Em: 454 – 516 nm. Additionally, a bright-field image was captured. The images were exported in the single channels or as overlays as 16-bit tagged image files after equal adjustment for gain and color intensity within one experiment.

## Acknowledgements

The work was supported by grants from the Deutsche Forschungsgemeinschaft SFB766/A7 and GRK1708 to CW. We thank Isabell Samp, Natalya Korn and Vittoria Bisanzio for excellent technical assistance. We thank Markus Bischoff for fruitful discussions and Jan Liese for providing plasmid pJL78. The mutants NE386 and NE425 from the Nebraska library were obtained through the Network on Antimicrobial Resistance in *Staphylococcus aureus* (NARSA) program. The authors have no conflict of interest to declare.

